# Protocol for measuring endocrine disruptive effects on transcriptional bursting using single-molecule imaging in human breast cancer cells

**DOI:** 10.64898/2026.05.01.722245

**Authors:** Pelin Yasar, Christopher R. Day, Joseph Rodriguez

## Abstract

Transcriptional bursts regulate gene expression by altering burst size or burst frequency. Here, we present a protocol that integrates fixed-cell smFISH and live-cell single-molecule imaging to analyze estrogen-responsive transcriptional bursting of the TFF1 gene in human breast cancer cell lines. This workflow enables measurement of burst size, burst initiation, and active allele frequency to determine how endocrine disruptor chemicals modulate transcriptional bursting dynamics.

For complete details on the use and execution of this protocol, please refer to Day, Yasar et al.^1^

## OVERVIEW

To begin characterizing the effects of endocrine disruptor chemicals (EDCs) on transcriptional bursting, single-molecule RNA fluorescence in situ hybridization (smFISH) is recommended due to its speed and cost efficiency. This method uses multiple 18-22 bp probes labeled with fluorescent dyes^2^. Hybridization of these probes to target RNAs produces localized fluorescence signals that appear as bright spots, allowing transcription site intensity to be quantified as burst size (Fig. 1A). These transcription sites are identified by colocalization of exon and intron smFISH signals (Fig. 1B). However, intron probe design must account for gene length and position. 5’ introns in long genes may be spliced prior to detectable signal accumulation, precluding colocalization and causing transcription sites to be missed. Additionally, using smFISH to quantify shifts in the prevalence of active transcription sites across a population can provide meaningful insight into alterations in bursting kinetics.

**Figure 1.**
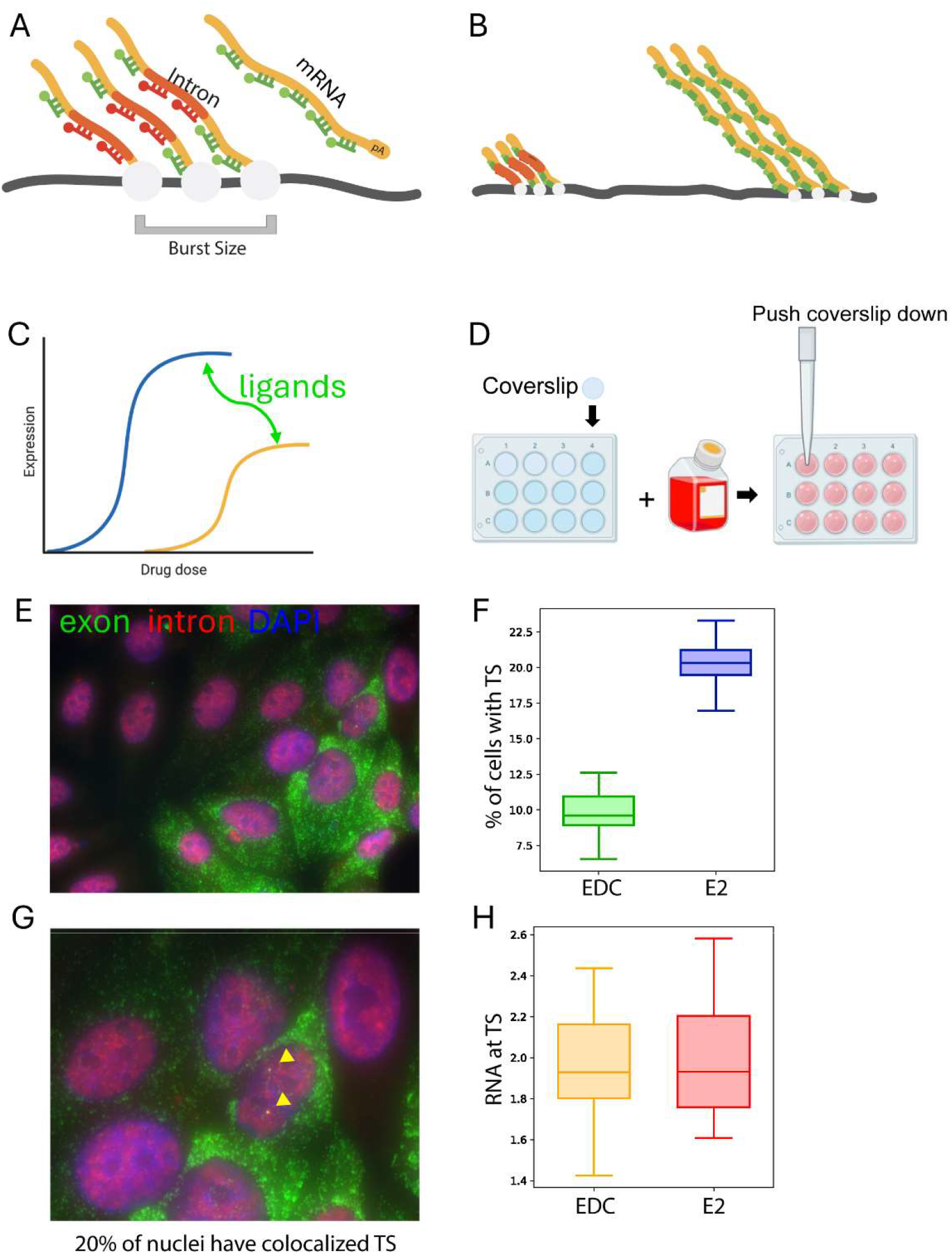
Quantification of transcriptional bursting using smFISH in response to ligand perturbations. (A) Schematic of smFISH-based detection of nascent and mature *TFF1* transcripts using exon- and intron-targeting probe sets. Transcription site (TS) intensity is used as a proxy of burst size. (B) Schematic of co-transcriptional splicing affecting intron probe detection. Early (5’) introns may be spliced before signal accumulation, leading to loss of intron signal at active transcription sites. (C) Schematic of ligand-dependent effects on transcriptional output. (D) Cell plating and coverslip preparation for smFISH imaging. (E) Representative smFISH image of *TFF1* transcripts. (F) Schematic illustrating calculation of the fraction of cells with transcription sites (% TS). (G) Magnified view of (E) highlighting transcription sites (TS). (H) Schematic illustrating estimation of RNA abundance at transcription sites, used to estimate burst size.

In this workflow, smFISH is first used to perform dose-response analysis and quantify the percentage of transcribing cells and transcription site intensities (Fig. 1C). Cells are plated on coverslips and prepared for imaging as described (Fig. 1D). These measurements allow quantification of transcriptional output and the fraction of responding cells across conditions (Fig. 1E-H). Based on these results, selected conditions are then examined using live-cell MS2 imaging to directly measure burst initiation and transcriptional dynamics.

Live-cell imaging provides complementary information on burst initiation and burst duration that cannot be obtained from fixed-cell methods^3^ (Fig. 2A, B). By combining these approaches, this protocol enables clearer mechanistic interpretation of how different ligands affect transcriptional bursting kinetics.

**Figure 2.**
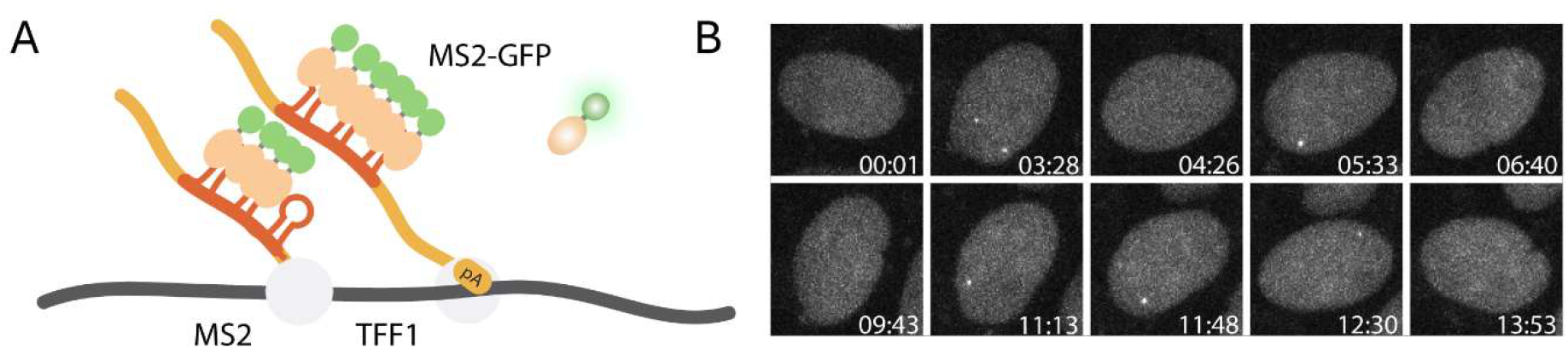
Measurement of transcriptional bursting dynamics using live-cell MS2 imaging. (A) Schematic of the TFF1-MS2 reporter system with MS2 stem loops integrated into the endogenous locus, enabling visualization of nascent transcripts via MS2-GFP. (B) Representative live-cell images showing transcription sites (TS) appearing and disappearing over time, consistent with bursting behavior.

## INNOVATION

Current descriptions of hormone-dependent gene regulation are largely based on bulk-cell measurements that average transcriptional outputs across populations, obscuring transcriptional bursting at individual loci. In contrast, single-cell studies demonstrate that only a subset of cells activates a given gene in response to hormone, revealing heterogeneous, burst-like transcriptional dynamics that bulk-based models fail to capture. Our protocol is innovative because it operationalizes a combined single-molecule RNA FISH (smFISH) and live-cell imaging workflow specifically designed to quantify transcriptional bursting in response to defined hormonal or ligand perturbations. smFISH provides rapid, high-throughput, and absolute measurements of RNA copy number and transcription-site intensity across large cell populations, enabling precise estimation of steady-state and burst-related parameters. In parallel, live-cell assays capture the timing, duration, and frequency of bursts at individual alleles, delivering longitudinal information that cannot be obtained from endpoint assays alone. By integrating these complementary datasets into shared kinetic models, this protocol constrains otherwise underdetermined parameters and links single-cell bursting features to population-level gene activation. Thus, the protocol provides a practical and reproducible framework to move from descriptive imaging to quantitative, mechanistic analysis of hormone-responsive transcriptional regulation.

### BEFORE YOU BEGIN

This protocol combines smFISH and live-cell MS2 imaging to analyze transcriptional bursting of the estrogen responsive TFF1 gene in human breast cancer cells (MCF7). The steps below describe the preparations required before beginning the experimental workflow.

Generate or obtain a *TFF1*-*MS2* reported cell line.

Live-cell imaging of transcriptional bursting requires a reporter cell line in which MS2 stem loops are integrated into endogenous *TFF1* locus.

1. Generate or obtain a *TFF1*-*MS2* knock-in cell line in which 24x MS2 stem loops are inserted into the 3’UTR of the TFF1 gene.
2. Express MS2 coat protein fused to GFP (MS2-GFP) in the reporter cell line to enable visualization of nascent transcripts.

**Note:** Generation of edited clones typically requires 3-6 months, depending on the cell line and genome editing strategy. Additional details on reporter construction are described in Rodriguez et al.^3^

Prepare hormone-depleted culture conditions.

To accurately measure ligand responses, cells should be pre-cultured in hormone-depleted medium before ligand treatment.

3. Culture MCF7 cells in phenol red-free medium supplemented with charcoal-stripped fetal bovine serum.
4. Perform hormone depletion for 2 days prior to ligand treatment to allow cells to reach steady-state transcriptional activity.

Order smFISH probe sets

Detection of transcription sites requires simultaneous hybridization of intronic and exonic RNA probes.

5. Order probe sets targeting exonic *TFF1* RNA labeled with one fluorophore (e.g., Quasar 570).
6. Order probe sets targeting intronic *TFF1* RNA labeled with a second fluorophore (e.g., Quasar 670) to enable two-color hybridization and transcription site identification.

Note: Probe set sequences for *TFF1* are provided in Table 1. Prepare hybridization and wash buffers

Prepare the smFISH hybridization and wash buffers described in the Materials and Equipment section prior to beginning the experiment.

### STEP BY STEP PROTOCOL

#### Dose Response smFISH^1^

Rationale: To ensure that observed effects are not due to suboptimal concentrations of ligand, a dose response of the *TFF1* bursting is recommended. The dose of maximal activity is used to compare against E2 maximal activity dose.

#### Seeding and Treatment

**Timing: 8-9 days**

1. Determine appropriate seeding density such that cells are 70-80% confluent on the day of fixation. For MCF7 cells which have a doubling rate of about 2 days, cells are seeded then allowed to recover for 2-3 days until the cell morphology is normal. This step is followed by 3 days of hormone depletion, and after addition of ligand addition are allowed to reach steady state for an additional 3 days. Therefore, the entire experiment takes 8-9 days.

**Note**: Plating MCF7 cells at 10-15% confluency is a good start. MCF7 cells have difficulty attaching to glass coverslips during the initial recovery.

2. Determine the number of samples to plate. For example, for 4 doses, plate three replicates for each dose which equals 12 samples.
3. Place 1 18mm No 1.5 coverslip into individual wells of a 12 well plate in the cell culture hood(Fig. 1D).
4. Add 1ml of complete media into each individual well.
5. Using a sterile pipette tip, push down on each coverslip to reduce probability of cell growth under coverslip.
6. Seed appropriate cell number into each well as determined in step 1 above.
7. Slowly move plate in up-down and left right directions to disperse cells. Avoid circular motions which can lead to a ring of cell growth or growth primarily in the middle of dish.
8. Allow cells to recover 2-3 days. Ensure cell morphology is normal.
9. Aspirate media and add 1ml of Hormone depleted media per well.
10. Return to incubator for 1hr.
11. Repeat steps 8-9 2X for a total of 3 washes.
12. After last wash, return cells to incubator for three days to allow cells to reach steady state.
13. Prepare 10 fold serial dilutions of ligand of choice in hormone depleted media.
14. Replace hormone depleted media on cells with dosed media.
15. Return to incubator for 3 days to allow cells to reach steady.

#### Cell fixation and permeabilization

**Timing: 1 h**

1. Wash cells 3x with 1ml PBS (1-3x)
2. Fix cells with fixation solution for 10 minutes at room temp (RT) in 4% PFA in PBS in fume hood.
3. Collect all PFA waste and washes. Do NOT aspirate.
4. Rinse cells 2X for 10 min with 1ml PBS.
5. Remove PBS and permeabilize fixed cells by incubating coverslips with 2ml 70% EtOH overnight at 4C.
6. Parafilm 12 well plate to reduce evaporation. Fixed cells can be stored and used up to 2 weeks if stored at 4C.

**Note**: An alternate permeabilization method which is immediately followed by smFISH hybridization is to incubate cells with 0.5% Triton PBS solution for 10 min. Rinse 1 x 10 min at 1x PBS. Then proceed to pre-hybridization step below.

#### smFISH Hybridization

**Timing: 6-7 h**

- Thaw Probe-Hybridization solution, probesets and smFISH wash solution
- Protect probesets from light exposure.

#### Prepare Probe-Hybridization solution

1. 50ul of probe mix is needed per 18mm coverslip. Add 2ul of diluted (1.25uM) probe stocks (2ul each for intronic and exonic) to 50ul of hybridization solution. Make 10% extra. This solution is very viscous.

**Note**: *The probe can be added directly from stock (12.5uM) or a 1:10 dilute working stock can be made.

2. Pre-hyb: Remove 12 well plate with samples from 4C and replace 70% ethanol with 1ml wash buffer and incubate for 2-5 min. Prepare hybridization chamber (step 3) while coverslips are in wash buffer.

**Note:** Prepare hybridization solution during this step.

3. For 12 samples, label the bottoms of two 10 cm petri dishes with the numbers 1-6, 7-12. These numbers should be evenly spaced out and correspond to the placement of the coverslip and enable tracking.
4. Critical Step: Cut two parafilm sections which will cover the inside of the petri dish. Cut the corners off with scissors. Remove wax paper and place parafilm, new side up. Using removed wax paper, push parafilm flush against petri dish bottom.

**Note**: Hybridizing on parafilm instead of directly on petri dish eliminates most bubbles which occur in probe hybridization solution during hybridization.

5. Place 50 ul of probe hybridization solution on each number.
6. While the coverslips are in wash buffer, use fine tweezers to gently lift the coverslip and grip the edge.
7. Gently dry coverslip edge one at a time using a kim wipe and place cell side DOWN onto probe mixture.

**Note**: Angled tweezers provide better maneuverability

**Note**: Do not remove wash buffer.

8. After all samples have been added to chambers, add a folded moist kim wipe (1ml of ddH20) to middle of each chamber. Add lid.

**Note**: Kim wipe keeps chamber humid

9. Seal chamber with parafilm and place in light protected 37 incubator for 4 hours at 37 C. Hybridization can also be performed overnight.

**Note**: Place a small beaker or tray of water in incubator to keep incubator humid.

#### Wash

10. At 3.5 hours, place wash buffer into 37 C incubator
11. Rinse coverslips with 1 ml wash buffer and incubate for 30min at 37 C.
12. Thaw Prolong gold with DAPI mounting media.
13. Rinse coverslips again with 1 ml wash buffer and incubate for 30min at 37 C.
14. Rinse quickly 1ml 1x with 2xSSC.
15. Rinse with 1ml 1x PBS and incubate for 5 min covered at room temperature.

#### Mounting

16. Briefly dry coverslips by placing against tip box on kim wipes. The goal is to remove excess liquid.
17. Add a very small drop of mounting media to glass slide by touching media drop to slide.
18. Gently place coverslip face down on mounting media. Cover with foil or place in drawer and let dry overnight.

**Note**: From this point on, it is critical to protect samples from light.

#### Microscopy

**Note**: Imaging can be performed on widefield microscopes. Key features to consider are light source and objective. The light source should have a powerful LED (Lumencor SpectraX) which can be found on custom setups. Additionally, it is important to ensure that the excitation and emission filters are optimal with the designed probesets. The objective magnification is also important as imaging at too low a magnification can lead to RNA spots that are located within single pixels. This makes it difficult to distinguish RNA spots from technical noise. 40X and 63X are good magnifications to use as single RNA spots are captured over several pixels.

1. To quantify transcription sites, a minimum of 1000-2000 cells is recommended. Using a 40X objective and the MCF7 cells at 70-80% confluency, a 5×5 grid is recommended.
2. The whole nuclear volume should be captured, so the number of Z-steps must be determined.

#### Transcription Site Analysis

The python pipeline in Day,Yasar et al^1^ is used for spot and cell segmentation. https://github.com/singlecelldynamics/AcuteBPA Pipeline will calculate max projections from 3D stacks of individual field of views. It will also identify spots in both channels, determine if the spots colocalize, and calculate the number of RNA at transcription sites. You will also need to have Cellprofiler^4^ (V. 3.1.9) installed to segment nuclei and cytoplasm. An IDE is recommended when working with this software. We use Spyder which can be installed with Anaconda.

1. After identifying transcription sites (TS) and quantifying burst sizes for all doses, compare % of Transcribing cells (%TS) and burst sizes at doses where activity plateaus.
2. Changes in burst size indicate that different reinitiation rates occur during a burst when cells are treated with the ligand relative to E2.
3. Changes in %TS when treated with the ligand, suggest one of at least two possibilities. Either the number of alleles that is active is different or the burst initiation is different.
4. To experimentally distinguish between these possible mechanisms, live cell imaging of the estrogen response is necessary.

#### Live Cell MS2 Imaging

The *TFF1*-*MS2* cell line for live cell imaging of the estrogen response was previously established^3^ by integrating 24X MS2 stem loops into the 3’UTR of the TFF1 gene (Fig. 2A, B).

Note: To capture transcriptional bursting dynamics at other genes or in different cell lines, 24X MS2 stem loops must be integrated into the endogenous locus using genome editing. Generation of single-cell edited clones typically requires 3-6 months depending on the cell lines^3^.

#### Seeding and treatment

1. Similar to seeding for the dose response smFISH, determine the appropriate number of cells to seed. The two chamber NUNC labtek dishes are appropriate
2. Plate TFF1-MS2 cells onto NUNC Labtek two chamber No. 1.5 glass dishes in complete media and allow the cells to recover for 2-3 days.
3. Confirm that morphology is normal.
4. Aspirate media and add 1ml of Hormone depleted media per well.
5. Return to incubator for 1hr.
6. Repeat steps 4-5 2X for a total of 3 washes.
7. After last wash, return cells to incubator for three days to allow cells to reach steady state.
8. Prepare specific dilutions of ligand of choice in hormone depleted media.
9. Replace hormone depleted media on cells with dosed media.
10. Return to incubator for 3 days to allow cells to reach steady.

#### Microscopy

Imaging can be performed on a widefield microscopy with laser excitation, Zeiss confocal with laser excitation or HILO system such as the Andor Dragonfly Spinning Disk Confocal/HILO Microscope.

**Note**: Excitation power must be optimized to reduce bleaching in nucleus.

**Note**: Cells with lower levels of coat protein (MS2-GFP) will provide better signal to noise. Brightest cells will have too much background diffusing MS2-GFP.

**Note**: Entire nuclear volume must be captured to ensure that when transcription site turns off, that it is not due to the transcription site moving out of the focal plane.

**Note**: Make sure you are using autofocus, otherwise focus will drift over the long imaging timescales.

**Note**: We normally use imaging intervals of 100 seconds which provides a good balance of time resolution and bleach prevention.

**Note**: To capture bursting dynamics, we image 512 frames at 100 second intervals and several Z stacks, overnight. This comes out to 14.2 hours of imaging. However, the exact time isn’t as important as being able to capture the changes in the inactive times. So for a gene that has an average inactive time of 1.5hrs, 8 hours would be enough. This could enable two imaging sessions per day.

**Note**: Depending on the speed of the stage, piezo and camera exposures, we maximize the number of region of interests (ROIs) that we can image in series. For example, we can image 9 field of views within 100 seconds on our setup.

1. Setup imaging of multiple tiles, and the entire nuclear volumes. Try 512 frames, every 100 seconds, 7 micron volume.
2. Perform a test run to see if fluorescence is bleaching rapidly. If so, reduce laser power and try again.
3. Aim to get at least 30-50 transcribing cells per condition. We generally only consider cells where the cell was observed in the movie the entire time.

### Analysis

#### Tracking cells and cropping

To extract transcriptional bursting dynamics from live-cell imaging data, a stepwise analysis pipeline is used, progressing from cell tracking to transcription site segmentation and kinetic modeling (Fig. 3).

**Figure 3.**
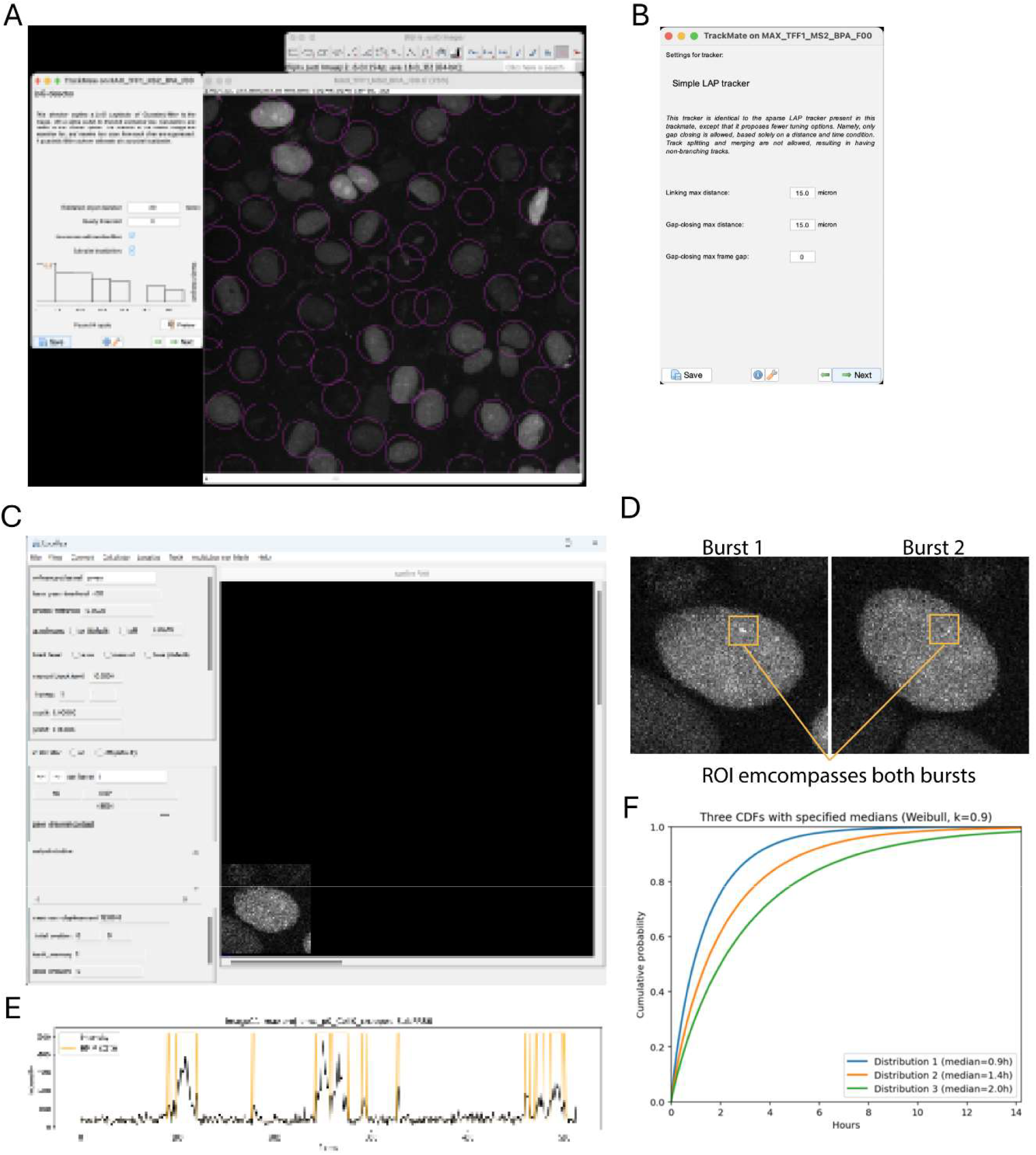
Analysis pipeline for transcriptional bursting from live-cell imaging. (A) Detection of nuclei using TrackMate with Laplacian of Gaussion (LoG) segmentation. (B) Configuration of LAP tracking parameters for linking nuclei across time. (C) Identification of transcription sites using Localize for spot detection and intensity extraction. (D) Region-of-Interest (ROI) selection encompassing consecutive bursting events from individual alleles. (E) Hidden Markov Model (HMM) fitting of transcriptional states to extract ON (active) and OFF (inactive) durations. (F) Schematic of cumulative distribution analysis of ON and OFF times to quantify burst duration and initiation frequency.

Over the course of a 10-14hr movie, cells move considerably in the field of view (FOV). Therefore, tracking and cropping of cells is important for MS2 spot segmentation and tracking. We accomplish this task with the use of the FIJI^5^, TrackMate^6^ plugin and python code.

#### FIJI

1. Calculate and save the maximum intensity projections of each FOV using FIJI. You should be left with a time series stack of each FOV.
2. Open a single TIFF stack in FIJI.
3. Select TrackMate from plugins→Tracking→TrackMate menu.
4. Enter initial parameters for the calibration settings if not set from image metadata, and select the next arrow button.
5. Select a detector, we use the LoG detector. Hit next arrow button.
6. Enter a nuclei diameter in pixels or microns for LoG detector. Press the Preview button to test how well the detector encases nuclei. Small circles will be drawn on the TFF stack (Fig. 3A). Too small, and it will find too many nuclei, too large and it can merge nuclei.
7. Press the next arrow button. It will identify regions in each time frame.
8. Continue pressing the next arrow button until you reach the “Select a Tracker”.
9. Select the LAP tracker and hit the next arrow button.
10. Enter linking distances, gap closing distances in the current window (Fig. 3B). For example use 15 microns for these parameters. Set gap-closing max frame gap to 0. We analyze only full length movies were the nuclei was identified in all frames. Press the next arrow button.
11. After TrackMate has made tracks, select the next arrow button twice.
12. Lines will appear over the tracked circles which illustrate the tracked circle’s path. Play the movie and evaluate tracking. If necessary go back and adjust parameters. Using this method, some cells may not be tracked for the entire movie.
13. A Display options window should be present. Select the Track button at the bottom. A table will appear with the tracked spots data. Save the data to a csv file for the FOV by pressing the Export to CSV button.

#### Switch to Python

14. Use **TrackMateTiffCrop.py** to crop cells using the track csv data. Enter paths, maximum intensity projections, and TrackMate csv files. Multiple files can be entered.
15. Enter a cropping window size. **TrackMateTiffCrop.py** will use this parameter to crop around the TrackMate csv file plus and minus that window parameter.
16. Cells should be cropped into individual tiff stacks. The first frame of the original maximum intensity projection will be saved along with the Cell IDs that were cropped. Use this as a guide to determine efficiency of cropping and to go back and extract remaining cells using other methods.

#### MS2 spot segmentation and tracking

The following section uses Localize^7^ for spot calling and segmentation.

**Note:** If a *TFF1* allele is not bursting in the first frame, Localize will not capture the first inactive duration. Therefore, movies must be prepared for each transcription site by adding the first instance of the transcription site to the beginning of the frame. This new movie is saved. After the spot is tracked, the first frame data is removed.

**Note:** There are two major steps in tracking alleles. Segmenting transcription spots and then linking close spots from adjacent frames together.

#### FIJI

1. Add first frame of the first burst at the beginning of the movie using FIJI Stacks→Tools→Substack. Alternatively, you can use **FirstFrameSubstack.ijm** FIJI macro. Save the movie.

#### Switch to Localize

**Note:** *TFF1*-*MS2* nuclei may rotate and dramatically change shape. We do not track cells that have changing shapes because of the uncertainty of allele organization.

Rotating nuclei maintain similar distances between multiple active alleles and although more difficult, are trackable. The easiest way to track allele bursting is by linking consecutive bursts of an allele. Then stitching together the output .**loc** file.

1. Run Localize (Fig. 3C). A user interface will appear. Open your TIFF stack. Scroll down to see it.
2. Enter the Following parameters:
  a. Auto bpass: **ON**
  b. frames: **1** to **Total Number of Frames**
  c. FCS mode: **ON**

**Note:** FCS mode ON is critical for analyses. With FCS mode ON, Localize will keep the last known position of the identified transcription site in the event it cannot find a transcription site in the current frame. It will extract intensity. This background intensity is critical for downstream analyses.

3. Determine the last frame of the next burst. Replace the **Total Number of Frames** in 2b **above** with this number.
4. From the menu, select Localize→segment with ROI. Draw ROI that encompasses two consecutive bursts from the same allele (Fig. 3D). In the initial case, the ROI will surround the first frame transcription site and the first actual burst.
5. After Localize completes, load the recently created .**loc** file. This is a text file with coordinates and intensities of all identified spots.
6. Set frame to first frame. You should see a box around the first spot. Check that if also found spots during your first burst. If not repeat with larger ROI.
7. Save .**loc** file with a unique name indicating frames.
8. Continue with first and second bursts of allele. Change frames in 2b accordingly and draw ROI’s to encompasses these two bursts.
9. Repeat until all bursts of allele have been tracked.
10. For the last burst, you will select an ROI that encompasses the last burst and create the last .**loc** file.
11. Combine all .**loc** files. Remove redundant overlapping frame data.
12. Load .**loc** file and select from menu, Track→Track. After tracking is complete, a .**trk** file is created. This file contains the coordinates of the tracked transcription site over time.

### Switch to FIJI

13. Check the tracked transcription site by creating an overlay of the track onto the original time series TIFF stack (Fig. S1).
14. In FIJI, load the **trkCheck.ijm** macro. Enter file and path information of the TIFF stack and .**trk** file into the macro. Run macro.
15. Play movie or advance through frames.

### Switch to HMM python script

1. It is recommended to shift the data and remove the first frame of the .**trk** file which is due to the
2. In an IDE such as Anaconda, load **traceHMMFit.py**. Enter path information for a directory with all .**trk** files belonging to same condition.
3. Run traceHMMFit.py. It will generate plots of the intensity traces **(.trk**) and the fit HMM states (Fig. 3E).
  a. Code will output median and mean ON and OFF (active and inactive) durations in minutes.
  b. Image files of the plots(.**hmm.png**). HMM state files ((.**hmm.txt)** also output for plotting in other software.
  c. The file **TrkONOFFSummary.txt** is created and contains the durations of each inactive and active period for all .**trk** files. There are two columns, State and duration.

**Note**: It is critical to inspect each HMM fit before accepting results.

4. Create cumulative distribution plots of the active and inactive distributions (Fig. 3F).

### Integrating smFISH and Live cell MS2 results

smFISH data can identify changes in burst size through the normalization of transcription site intensities and comparing this measurement between different conditions (Fig. 4). Additionally, changes in the percent of transcribing cells can indicate changes in active alleles across the population, or changes in burst initiation. To distinguish between these mechanisms, using live cell data is critical as burst initiation is directly measured. If no changes in burst initiation are observed, the interpretation of the smFISH data is that fewer alleles are in active states as previously observed^1^. Live cell MS2 imaging also provides data on burst duration. This measurement is the dwell time of the MS2 labeled transcript and the dwell time is the sum of several processes including burst size, elongation rates, termination and transcript retention. Pairing changes in active duration with changes in burst size can help clarify the mechanistic interpretation.

## EXPECTED OUTCOMES

smFISH assays using exon and intron probes targeting the TFF1 gene are expected to yield bright, punctate RNA spots spanning several pixels, with line-scan intensity profiles showing a single, centered peak above background. Intron-labeled spots should be localized predominantly to the nucleus, whereas exon-labeled spots should be detected in both the cytoplasm and nucleus. Nuclear colocalization of exon and intron spots will mark active transcription sites (TSs), and individual aneuploid MCF7 cells should exhibit 0-5 TSs under typical induction conditions. From these images, users should obtain heterogeneous RNA per cell distributions that are visually apparent in the raw images and quantifiable across at least ∼1000 cells per condition. Under saturating estradiol stimulation, *TFF1* expression is expected to reach on the order of ∼70 RNA molecules per cell (cell line-dependent), and boxplot analysis of RNA at colocalized TSs should yield median burst sizes of approximately 1.5-3 RNA per TS for E2 doses. Comparing RNA per cell, fraction of cells with ≥ 1 TS, and RNA per TS across ligands or perturbations will reveal how total transcriptional output, fraction of responding cells, and burst size change between conditions.

Live-cell MS2 imaging in the *TFF1*-MS2 reporter line is expected to produce nuclei with a broad range of MS2-GFP expression levels; 20-30% of cells may lack detectable MS2-GFP signal and will not display visible transcription sites. Cells with excessively high MS2-GFP expression will show elevated nuclear background that precludes clear visualization of TSs. In optimally expressing cells, 1-3 bright punctate TSs, clearly above background, should be visible in a minority of nuclei. These TSs will appear for multiple consecutive frames, disappear for variable intervals, and then reappear, consistent with transcriptional bursting. Some cells without initial TSs but with adequate MS2-GFP levels may show burst onset at later time points during acquisition. For quantitative analysis, users should obtain ≥ 30 high-quality intensity traces (tracked TSs) per condition, enabling construction of ON- and OFF-time histograms and their corresponding cumulative distributions. Comparing cumulative ON- and OFF-time distributions across ligands or conditions will indicate changes in burst duration (ON time) and burst frequency or initiation (OFF time).

Integrating smFISH and MS2 outcomes will allow mechanistic interpretation of how specific ligands or perturbations modulate *TFF1* transcriptional bursting. For example, E2 dose response may primarily alter burst frequency, increasing both the fraction of cells with TSs and the number of bursts per allele, whereas TRIM24 inhibition may reduce burst size with minimal effects on the fraction of active cells^3,8^. Changes in ON-time distributions without changes in burst size would be consistent with altered burst duration (e.g., following enhancer perturbation), whereas changes in OFF-time distributions would support modulation of burst initiation frequency. Together, these combined readouts enable users to distinguish changes in burst frequency, burst size, burst duration, and number of active alleles, rather than attributing all effects to overall RNA output.

## LIMITATIONS

The present protocol quantifies transcriptional bursting at a single endogenous locus, *TFF1*, and evaluates how ligands or perturbations modulate its bursting dynamics. Extending this combined smFISH and live-cell MS2 framework to additional genes would require generating new cell lines with MS2 stem loops integrated at each target locus, which is technically demanding and time consuming, and thus may limit throughput and generalizability across loci. Measurements of burst duration from MS2 imaging also represent the total dwell time of the nascent, MS2-labeled transcript on chromatin and therefore convolve polymerase elongation through the 3′ MS2 cassette, polyadenylation, and termination. As a result, changes in the observed ON time cannot be unambiguously attributed to a specific kinetic step without additional, locus-specific controls or complementary assays.

## TROUBLESHOOTING

### Problem 1

Transcription sites are not visible or nuclear background is too high in MS2 imaging

### Potential solution

MS2 coat protein (MCP-FP) expression is likely too high, causing diffuse nuclear fluorescence that obscures puncta. Reduce MCP-FP expression and use flow cytometry to select a subpopulation with intermediate nuclear signal, excluding very bright and very dim cells, so that nuclei are clearly labeled but 1-3 bright transcription sites remain distinguishable.

### Problem 2

No transcription sites are visible and nuclei are poorly labeled in MS2 imaging

### Potential solution

MS2 coat protein expression is likely too low to label MS2-tagged RNAs and nuclei efficiently. Adjust the expression cassette (for example, promoter strength or selection strategy) and repeat flow-based enrichment to obtain cells with moderate nuclear fluorescence, verifying suitable signal with short test movies before long time-lapse acquisition.

### Problem 3

Weak or no smFISH signal

### Potential solution

Hybridization conditions are likely suboptimal. Optimize hybridization temperature in a narrow range around the recommended value and adjust formamide concentration to maximize punctate spot intensity while minimizing background, testing a small matrix of conditions for each new probe set.

### Problem 4

Poor probe penetration or inconsistent smFISH signal across samples

### Potential solution

Fixation/permeabilization may not be optimal for the sample type. If paraformaldehyde plus ethanol permeabilization yields weak or heterogeneous signal, test paraformaldehyde fixation followed by brief permeabilization with Triton X-100, titrating detergent concentration and incubation time to improve probe entry while preserving morphology.

### Problem 5

No signal from far-red smFISH probes

### Potential solution

The imaging system may not be properly configured for the far-red dye. Confirm that the microscope has an appropriate excitation source and filter set for your far-red fluorophore (for example, a Cy5- or Alexa Fluor 647-compatible filter set) and validate with a positive-control sample or fluorescent beads in the same spectral range.

## RESOUCE AVAILABILITY

### Lead Contact

Further information and requests for resources and reagents should be directed to and will be fulfilled by the lead contact, Joseph Rodriguez.

## AUTHOR CONTRIBUTIONS

P.Y., C.R.D., and J.R. conceived the project. P.Y., C.R.D., and J.R. wrote the original draft and reviewed and edited the manuscript.

## ACKNOWLEDGEMENTS

We thank the members of the Rodriguez Laboratory and the NIEHS Epigenetics and RNA Biology Laboratory for ongoing support, insightful discussions, and constructive feedback. We thank Jeff Tucker and Erica Scappini of the NIEHS Fluorescence Microscopy and Imaging Center for technical assistance with imaging. This research was supported by the Intramural Research Program of the NIH, National Institute of Environmental Health Sciences (ES103331 to J.R.). The contributions of the NIH author(s) are considered Works of the United States Government. The findings and conclusions presented in this paper are those of the author(s) and do not necessarily reflect the views of the NIH or the U.S. Department of Health and Human Services.

