## Supplementary Figures for "Protocol for measuring endocrine disruptive effects on transcriptional bursting using single-molecule imaging in human breast cancer cells"

**SUPPLEMENTAL FIGURE 1**

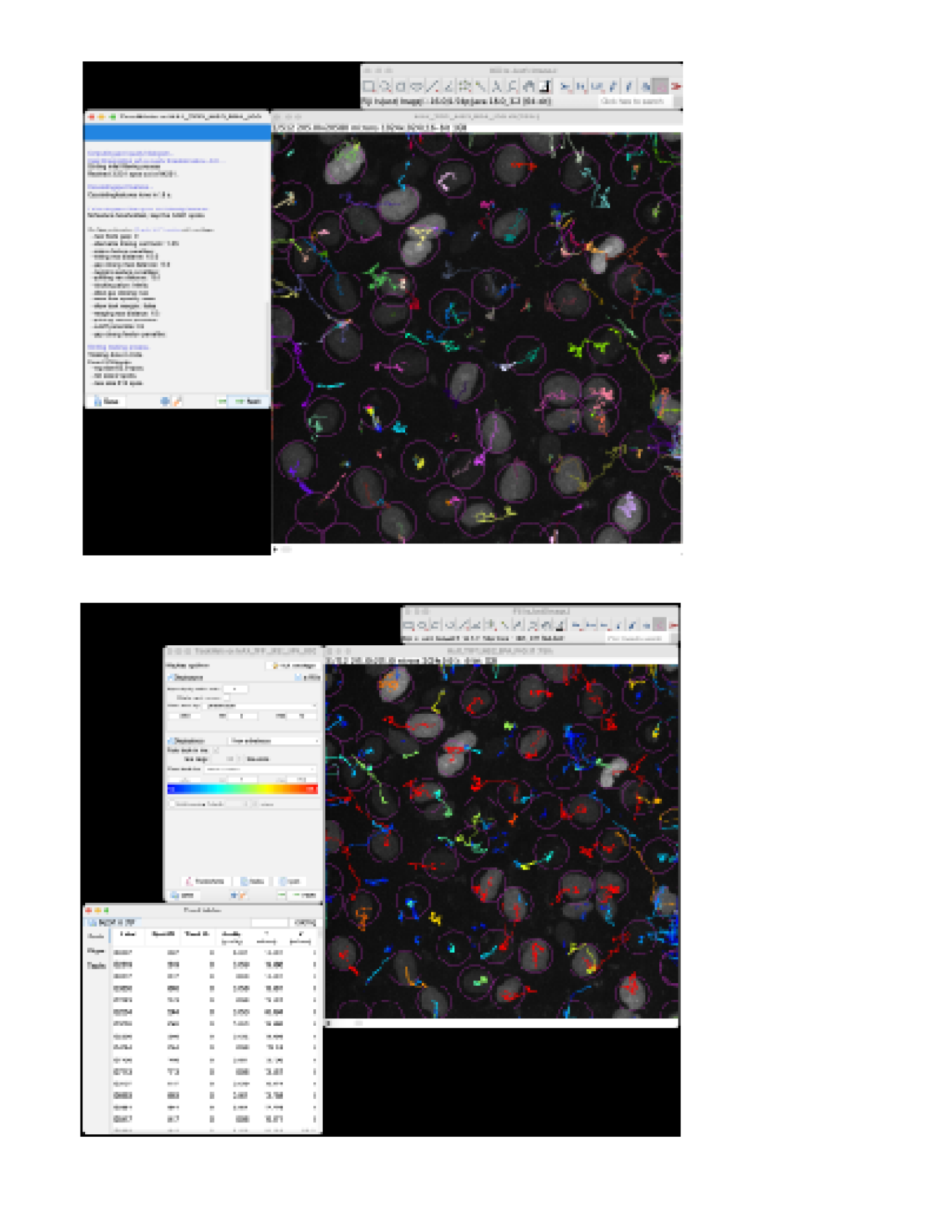

Figure S1 Validation of nuclei detection and tracking using TrackMate

(A) Detection of nuclei using LoG segmentation in TrackMate, with outlines overlaid on the original image.
(B) Tracking output showing linked trajectories across time, with individual nuclei assigned distinct tracks and corresponding tracking statistics.
